# QuickFit: A high-throughput RT-qPCR-based assay to quantify viral growth and fitness *in vitro*

**DOI:** 10.1101/2024.07.08.602587

**Authors:** Nicolas M.S. Galvez, Maegan L. Sheehan, Allen Z. Lin, Yi Cao, Evan C. Lam, Abigail M. Jackson, Alejandro B. Balazs

## Abstract

The quantification of viral growth rates is key to understanding evolutionary dynamics and the potential for mutants to escape antiviral drugs. Defining evolutionary escape paths and their impact on viral fitness allows for the development of drugs that are resistant to escape. In the case of HIV, combination antiretroviral therapy can successfully prevent or treat infection, but it relies on strict adherence to prevent escape. Here, we present a method that enables the quantification of viral fitness termed QuickFit, which employs large numbers of parallel viral cultures to accurately measure growth rates. QuickFit consistently recapitulated HIV growth measurements obtained by traditional approaches but with significantly higher throughput and lower error. This method represents a promising tool for rapid and consistent evaluation of viral fitness.

## INTRODUCTION

Understanding viral fitness and how treatments apply selection pressure on viral populations is pivotal for the development of drugs that prevent or treat infectious diseases (1). Viral fitness can be assessed at either the level of individual host infectivity or viral spread at the population level (2). Changes in the fitness of many viruses causing diseases in humans, such as HIV, Influenza, and SARS-CoV-2, have been extensively reported using *in vitro* and *in vivo* models to study drug resistance and immunological escape (1–3). Most *in vitro* approaches consider the outgrowth of viruses in cell cultures, followed by the measurement of either genetic material or viral antigens. Assays based on quantifying the viral genetic material are typically more reliable than those quantifying viral proteins, but are more expensive to run (4–6). However, the intrinsic measurement error of current technologies coupled with the stochasticity of viral infection and replication results in significant uncertainty in existing measurements of viral growth.

Despite significant public health initiatives and research efforts to develop therapies and vaccines, HIV remains a significant global pandemic (7). Highly active antiretroviral therapy (HAART) has been used for the past three decades to treat or prevent HIV infection (8–10), and several drug classes have been developed targeting key steps in the viral life cycle, including reverse transcription (RT), integration, and viral particle maturation (11–13). Most regimens combine two or three RT inhibitors, and strict adherence to the prescribed drug regimen is essential, as viral load rebounds within weeks of treatment interruption (10). As such, the development of long-lasting and curative strategies for HIV remains a priority for the field (14).

It has been extensively reported that suboptimal concentrations of early generations of HAART lead to the natural emergence of mutations in the RT polymerase, rendering the virus resistant to drugs (15–17). However, these resistance-associated mutations usually incur a cost in terms of viral replication rate, a phenomenon usually referred to as fitness cost (18–20). Determining accurate viral fitness of escape mutants is key to developing combinations of drugs that can effectively apply selection pressure on the virus, as mutations incurring higher fitness costs are less likely to arise (20). Existing approaches to determine viral growth and fitness of HIV and other viruses rely on either culture of the virus in cells *in vitro* or in animals (21). Longitudinal samples are subjected to either ELISA- or RT-qPCR-based methods to measure viral antigens or genetic material, respectively (22). However, these approaches are time-consuming, expensive, and have varying degrees of sensitivity (23,24), which leads to inconsistent reports of growth rates in the literature (25–27).

To address these limitations, we developed a high throughput, low-cost pipeline that could be used to systematically study viral fitness across different research centers. We developed an approach that was agnostic to differences in viral titers between stocks and that took advantage of improvements in high-throughput liquid handling to minimize hands-on time and maximize useful data. We employed RT-qPCR, given its wide dynamic range and high accessibility, to conduct an assay we termed QuickFit. To validate the method, we evaluated the relative growth rate of different HIV strains and compared them to those generated by a widely used p24-based ELISA. For each HIV strain tested, QuickFit generated robust and reproducible estimates of viral growth with higher precision than ELISA. To evaluate the fitness cost of drug escape mutations within the HIV RT *polymerase* gene, we measured the growth rates of viruses harboring known RT mutations. QuickFit was also used to determine the fitness cost imposed on viruses grown in the presence of Emtricitabine and the doubling time of numerous HIV isolates, finding significant differences in their growth rates.

## RESULTS

### Parallel viral cultures enable accurate modeling of growth rates

CD4^+^ T cells were isolated from peripheral blood mononuclear cells (PBMCs) using immunomagnetic purification and activated via CD3/CD28 co-stimulation to maximize their susceptibility to viral infection. To reduce experimental noise, we employed 16 parallel cultures for each individual virus under evaluation. Cells were mixed with serial dilutions of each virus and cultured over a period of 6 days to allow for multiple rounds of infection and viral replication. At daily intervals, beginning at 2 days post-infection, virus-containing supernatant was collected from each individual well simultaneously using a liquid handler and was immediately subjected to RNA extraction using a commercially available QuickExtract buffer and stored at -80°C in a 384-well plate format. After six days of growth, we determined viral loads for each sample by performing high-throughput RT-qPCR with strain-specific primers. Viral growth rates were determined by non-linear mixed-effects (NLME) modeling simulations in the MonolixSuite software using a half-maximal equation (**Figure 1**).

**Figure 1.**
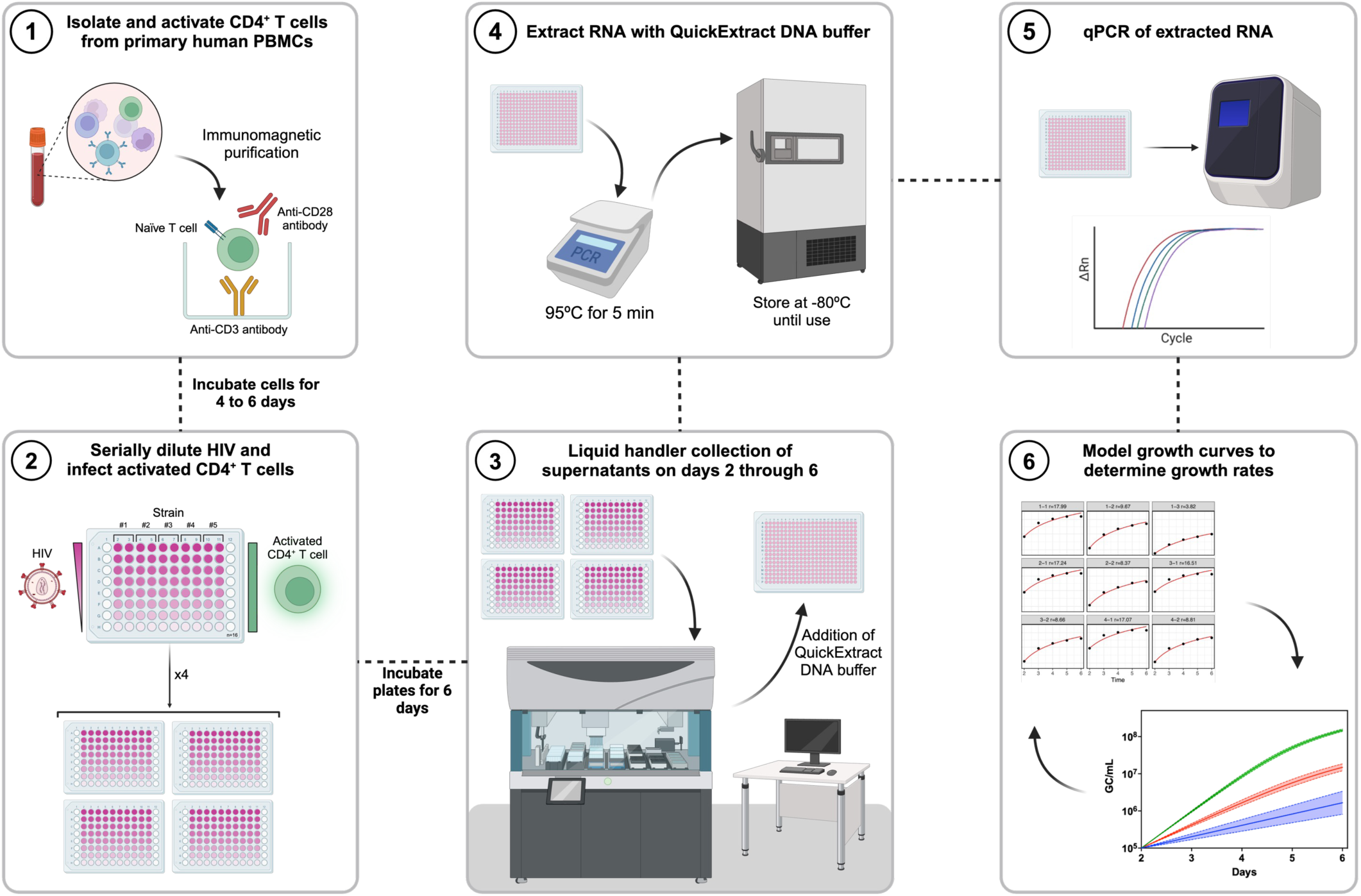
QuickFit is a high-throughput RT-qPCR-based HIV *in vitro* replication assay platform. Schematic representation of the experimental setup followed to perform QuickFit RT-qPCR evaluations of HIV growth rates *in vitro*.

To assess the robustness of QuickFit relative to the existing p24 ELISA-based methods, we infected activated CD4^+^ T cells with serial dilutions of HIV_REJO.c_, a clade B, R5-tropic, transmitted/founder (T/F) isolate that is widely used both *in vitro* and *in vivo* (28,29). Culture supernatants were collected daily and split into two samples that were either lysed to solubilize p24 protein or RNA extracted to quantify virus via p24 ELISA or RT-qPCR, respectively. Of the 16 samples evaluated via ELISA, only samples from cultures receiving the highest inoculum were above the limit of detection for the initial time point (**Figure 2A**). By day 4, the samples with the most and second-to-most diluted virus inoculum remained below the limit of detection (**Table S1**). In contrast, samples obtained in parallel that were quantified via RT-qPCR were within the limit of detection of the assay, irrespective of the time point evaluated (**Figure 2B**).

**Figure 2.**
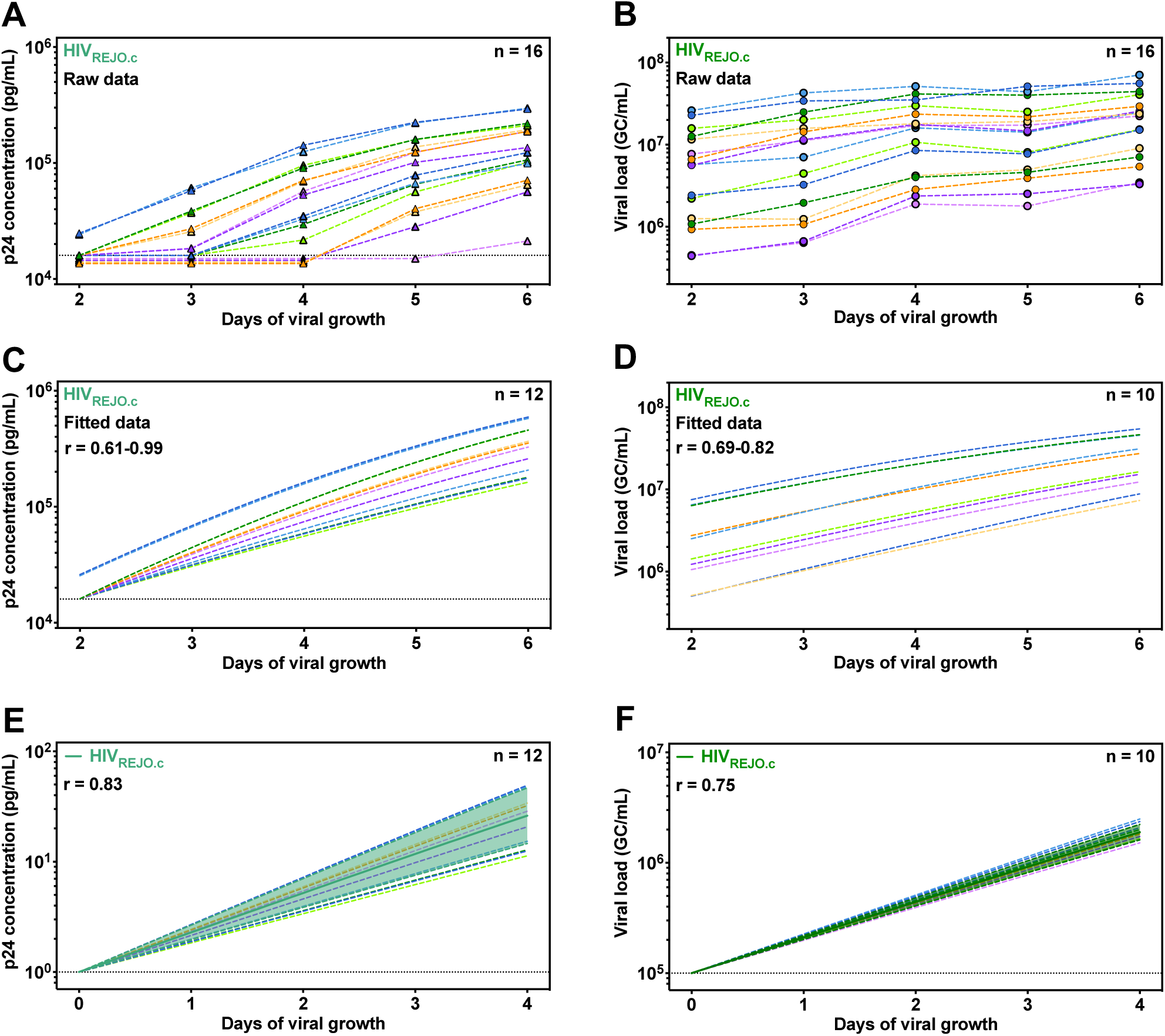
Analysis and modeling of HIV *in vitro* replication growth curves as measured by p24 ELISA and QuickFit RT-qPCR. Both p24 ELISA (**left**) and QuickFit RT-qPCR (**right**) were performed with samples obtained from HIV_REJO.c_ infected CD4^+^ T cells. (**A**) Determined raw p24 concentrations (pg/mL) and (**B**) viral loads (GC/mL) are shown. A total of 16 samples at 5 different time points were evaluated, and matching light-dark colors represent duplicates. (**C,D**) Samples were subjected to inclusion and exclusion criteria, and the filtered data was used to determine growth rates and carrying capacities by running an NMLE modeling with a half-maximal equation in the MonolixSuite 2021R software. (**E,F**) Viral growth (p24 concentration or GC/mL) was calculated by normalizing the initial value on day 0 to 1.00 pg/mL or 1×10^5^ GC/mL and subsequently interpolating the remaining values from days 1 to 4 based on the fit curve model. Overlaid is the mean growth rate and the 95% confidence interval. The r value represents the average growth rate determined for all included samples by each assay. Dotted lines represent the limit of detection for each assay.

We first analyzed the p24 ELISA data using a NLME model but found that including the concentrations from all wells yielded inaccurate growth estimates, as many of these were below the limit of detection at early time points. To address this limitation, we defined inclusion and exclusion criteria to enable consistent data analysis across independent *in vitro* experiments. We included all of the individual wells that were above the limit of detection after four days of viral growth, but excluded wells that reached the upper limit of detection at any time point. In addition, we excluded wells in which signals declined over 10% between any of the time points to remove wells exhibiting inconsistent growth. Together, these filters resulted in the exclusion of four individual wells (**Table S1**). NLME modeling of the 12 remaining wells via ELISA data generated growth rates (r) ranging from 0.61 to 0.99 (**Figure 2C**).

In contrast to ELISA measurements, RT-qPCR-based viral loads obtained from RNA-extracted samples were all within the limit of detection at all timepoints. When analyzing the viral load data from all wells, NLME modeling generated r values ranging from 0.62 to 1.06 (**Figure S1A and Table S2**). To increase the reliability of the analysis, we excluded wells that reached the upper or lower limit of detection at any time point, and wells in which the viral load declined by more than 10% between consecutive time points. To avoid measurements close to the carrying capacity of the assay, we excluded wells in which the viral load at the first time point was within 4-fold of the highest viral load observed. To avoid measurements exhibiting growth outside of the linear range, we also excluded wells in which the viral load at the last time point was more than 40-fold higher than the initial time point. This resulted in the exclusion of six individual wells (**Table S1**), and NLME modeling of the remaining wells yielded r values ranging from 0.69 to 0.82 (**Figure 2D**). Both p24 and qPCR-based NLME modeling were fit to a shared carrying capacity to enable direct comparisons of the two assays.

To visualize the average r value and error in these measurements, we simulated viral growth *in silico* starting from either a 1 pg/mL p24 concentration or 1×10^5^ GC/mL viral load using the previously determined growth rates. The average r value for the 12 individual wells analyzed by ELISA was 0.83, with a standard deviation of 0.12 (**Figure 2E and Table S2**). For qPCR data, the average r value across all 16 wells was 0.80, with a standard deviation of 0.12 (**Figure S1B, S1C and Table S2**), whereas after excluding unreliable wells, the average r value across the 10 remaining wells was 0.75, with a standard deviation of 0.04 (**Figure 2F, S1D, and Table S2**). Overall, the growth rates determined for HIV_REJO.c_ by QuickFit were similar to those obtained by ELISA but with significantly less variability and substantially less hands-on time, highlighting the advantages of QuickFit over conventional methods to evaluate viral growth.

### Comparison of HIV_REJO.c_ mutant growth rates via QuickFit

After validating QuickFit using a wildtype virus, we sought to measure differences in the fitness costs of individual mutations within HIV_REJO.c_ polymerase. Mutations M184I and M184V, which spontaneously emerge in people living with HIV who receive early generations of HAART, lead to drug resistance at the expense of fitness, by reducing reverse transcriptase processivity (30,31). Prior studies have described a range of impacts on viral replication kinetics and titer when comparing WT and Pol-mutant strains (31,32). We introduced Pol M184I and M184V mutations into the HIV_REJO.c_ infectious molecular clone (IMC) and prepared working viral stocks. We infected activated CD4^+^ T cells with WT or Pol mutant strains and collected supernatants to perform p24 ELISA and RT-qPCR. As these mutants have been shown to confer resistance to the antiretroviral drug Emtricitabine (FTC), we also included wells treated with 70 ng/mL (300 nM) of FTC, the equivalent of 0.5x times the EC_50_ value reported for this drug *in vitro* (33). Control samples were given the same volume of DMSO diluent without FTC.

Since these mutants shared the same parental strain, we performed NLME modeling using a shared carrying capacity. The average RT-qPCR r value generated for the WT HIV_REJO.c_ was 0.85 (± 0.09) (**Figure 3**). As previously reported, both *pol* mutations resulted in decreases in the r value, which translated into slower-replicating viruses, even in the absence of drugs. To quantify these differences, we normalized the r values to that of the WT strain to determine a fitness cost metric. Relative to the WT strain, the M184I had a fitness cost value of 1.31 (i.e., a 31% slower replication rate), whereas the M184V mutation had a fitness cost value of 1.55 (**Figure 3 and Table S3)**.

**Figure 3.**
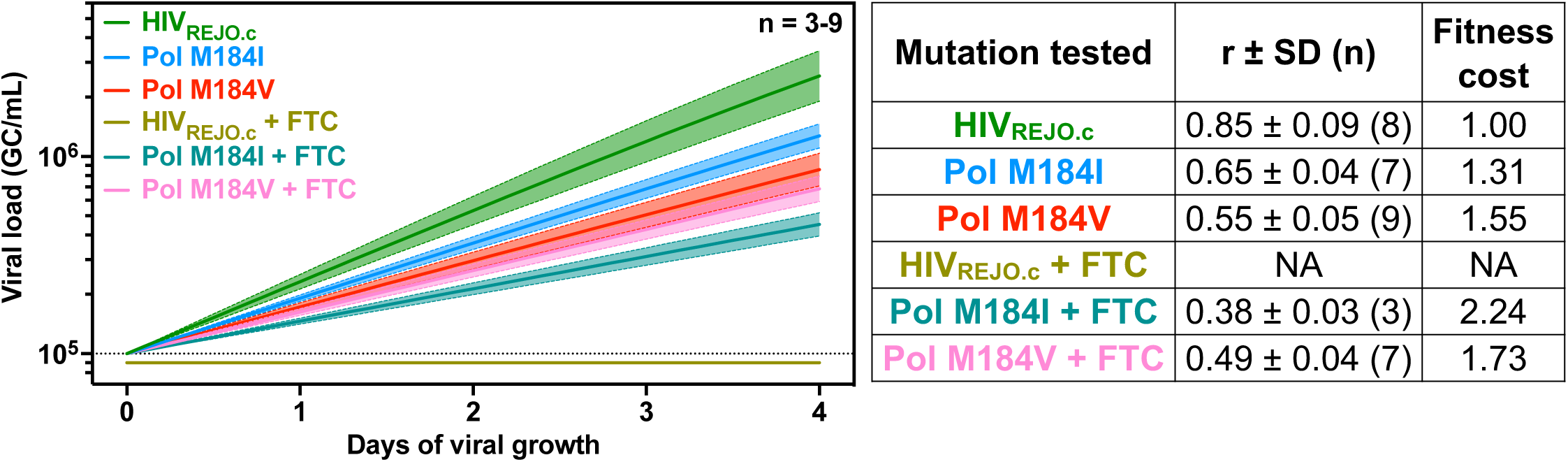
Comparative assessment of replication rates among HIV_REJO.c_ mutants for the *pol* gene in the presence of antiretroviral drugs. Single-point mutants for the *pol* gene were generated using the HIV_REJO.c_ IMC backbone. Then, QuickFit RT-qPCR was performed, and growth rates were determined. To evaluate the impact of ART drugs in these mutants, these strains were also grown in the presence of 300nM of emtricitabine (FTC, 0.5x the EC_50_ *in vitro* value) or DMSO as a control. Viral growth (GC/mL) was calculated by normalizing the initial value of each sample (after inclusion and exclusion criteria filtering) on day 0 to 1×10^5^ GC/mL and subsequently interpolating the remaining values from days 1 to 4 based on the fit curve model. Overlaid is the mean growth rate and the 95% confidence interval. The table to the right shows the r values (growth rates) ± the standard deviation for each strain tested for both conditions. The sample size (n) references the number of wells included in the analysis after filtering with inclusion and exclusion criteria. The fitness cost represents the doubling rate of each strain relative to the WT HIV_REJO.c_ without FTC. The dotted line represent the limit of detection for each assay. NA: No growth was detected.

In contrast to the DMSO-treated controls (**Figure S2A,B**), treatment with FTC resulted in no growth for the WT HIV_REJO.c_, as measured by both p24 ELISA and RT-qPCR (**Figure S2C,D**, top). However, both *pol* mutants were able to grow in the presence of FTC to varying degrees (**Figure S2C,D**, middle and bottom). Fitness cost values for M184I and M184V in the presence of FTC were 2.24 and 1.73, respectively. We found similar r values using data generated by ELISA, but with significantly greater variability which masked differences between each condition evaluated (**Figure S2E and Table S3**).

Sources of variability when evaluating HIV growth and fitness include both intrinsic donor PBMC differences as well as degree of cellular activation between experiments. To address the robustness of QuickFit across donors, we infected activated CD4^+^ T cells from three different individuals in three independent experiments with HIV_REJO.c_ and the Pol M184I strain (**Figure S3**). No statistical differences were seen between any of the three donors tested when comparing the growth rates for HIV_REJO.c_ WT or the Pol M184I strain (**Figure S3A**). Accordingly, growth curves for each strain were within the measurement error for all donors (**Figure S3B**), and pairwise comparisons found that fitness of the M184I mutant relative to wildtype was very similar, irrespective of the donor (**Figure S3C**). These data suggest that donor-to-donor variability does not significantly impact QuickFit when comparing mutants of the same strain.

Finally, to confirm that growth rates determined for FTC-treated samples were not impacted by the presence of RT inhibitor drug in the culture supernatants, we performed RT-qPCRs with serial dilutions of previously titered HIV_REJO.c_ viral stocks spiked with increasing concentrations of FTC (ranging from 35 ng/mL (60nM) to 700 ng/mL (30µM)). No differences were observed for any of the raw CT values across different viral dilutions or FTC concentrations (**Figure S4**). Accordingly, CT values reported in the presence of FTC or DMSO were the same as those of non-spiked viral stocks. Altogether, these results demonstrate that QuickFit RT-qPCR can be used to reliably determine growth rates for mutant strains of the same HIV isolate.

### The high sensitivity of QuickFit allows for the evaluation of different HIV isolates

To quantify differences in HIV growth rates between isolates, we tested four different HIV strains: HIV_NL4-3_ a clade B, X4-tropic, chronic isolate; HIV_JR-CSF_ a clade B, R5-tropic, chronic isolate; HIV_89.6_ a clade B, dual-tropic (X4- and R5-tropic), chronic isolate; and HIV_BF520_ a clade A, R5-tropic, T/F isolate (BEI Resources, #ARP114, #ARP2708, ARP#3552, #ARP13402, respectively) (34,35). All isolates could be detected by p24 ELISA and RT-qPCR, albeit with differences in sensitivity (**Figure S5**). Robust detection of p24 was observed for HIV_NL4-3_, HIV_JR-CSF_, and HIV_89.6_ (**Figure S5A,C,E**). However, HIV_BF520_ was difficult to detect, with only the two wells with the highest initial input of the virus reaching values above the limit of detection by day 2 (**Figure S5G**). In contrast, RT-qPCR-based viral loads were all above the limit of detection for HIV_NL4-3_ and HIV_89.6_, while 14 out of the 16 samples for HIV_BF520_ and HIV_JR-CSF_ were detectable at the first time point (**Figure S5B,D,F,H**).

We modeled each virus growth rate and carrying capacity independently, finding that average r values and standard deviations determined by either ELISA or RT-qPCR were similar for the four strains (**Figure 4 and Table S4**). To enable direct comparison between strains, we performed a single simulation including data from all strains without constraining the carrying capacity to a shared value and the relative growth rates were expressed as doubling time. All strains had a doubling time of approximately 24 hours, in line with previous reports (36) (**Figure 5 and Table S5**), with HIV_REJO.c_ and HIV_JR-CSF_ exhibiting the largest difference in growth rates. Finally, to determine whether the growth rates of different HIV isolates are impacted by donor-to-donor differences, PBMCs from three different donors were infected in three independent experiments with either HIV_REJO.c_ or HIV_JR-CSF_. A single simulation including both strains and all donors was performed without constraining the carrying capacity to a shared value. Interestingly, growth rates measured in cells from donor 1 were significantly faster than those determined for Donors 2 and 3 (**Figure S6A**). However, no differences were seen across donors when relative fitness was measured by normalizing growth rates (**Figure S6B**). Taken together, our results demonstrate that QuickFit can be used to evaluate and compare the growth rates, carrying capacities, and fitness of different HIV strains with high precision.

**Figure 4.**
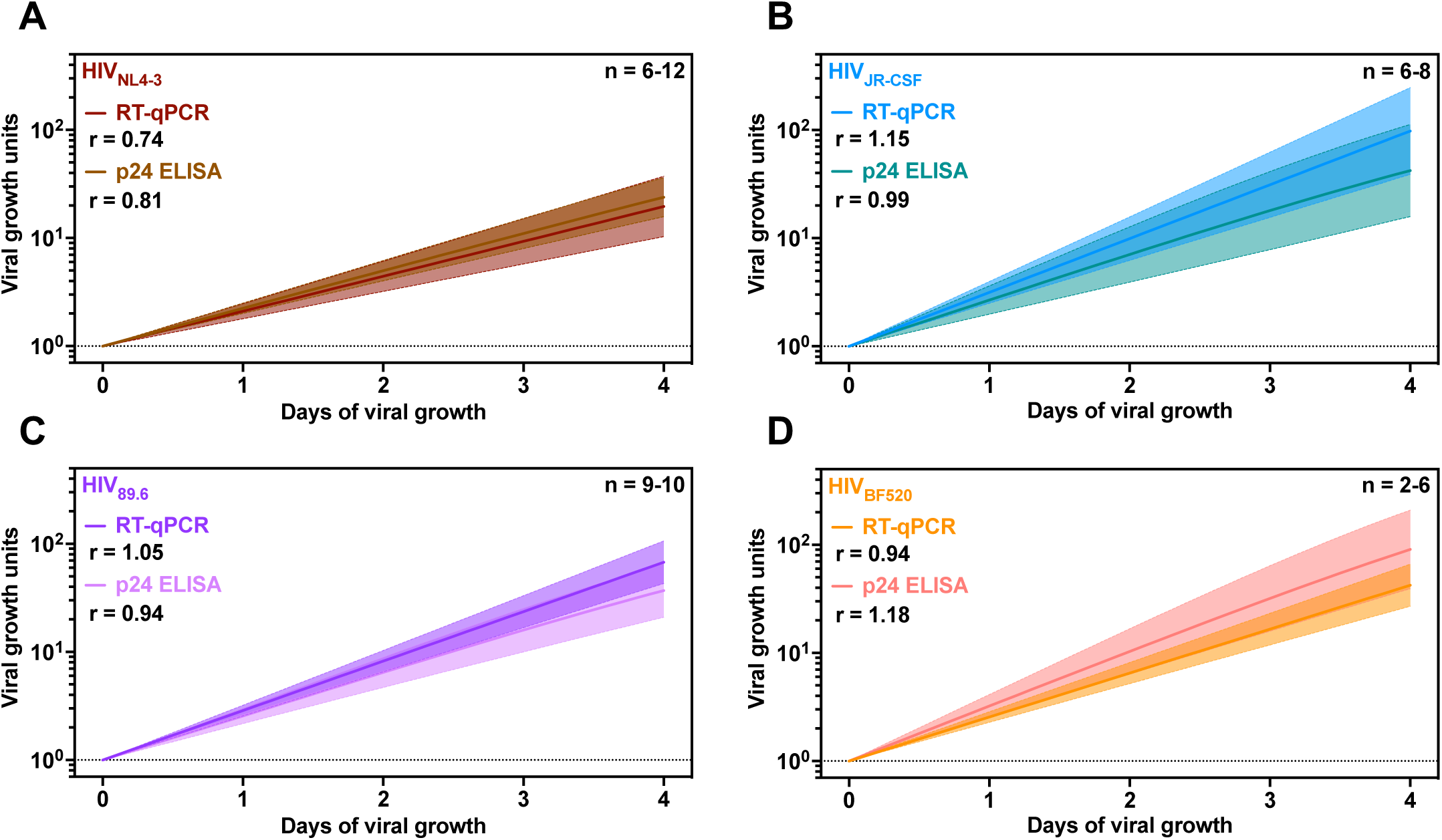
Evaluation of viral growth of four different HIV-1 isolates by p24 ELISA and QuickFit RT-qPCR. (A) HIV_NL4-3_, (B) HIV_JR-CSF_, (C) HIV_89.6_, and (D) HIV_BF520_ growth rates were determined using the QuickFit pipeline by p24 ELISA (lighter shades) and RT-qPCR (darker shades). Viral growth (p24 concentration or GC/mL) was calculated by normalizing the initial value on day 0 to either 1.00 pg/mL or 1.00 GC/mL and subsequently interpolating the remaining values (filtered by inclusion and exclusion criteria) from days 1 to 4 based on the fit curve model. Overlaid is the mean growth rate and the 95% confidence interval. The r value represents the average growth rate determined for all included samples by each assay. Dotted lines represent the limit of detection for each assay.

**Figure 5.**
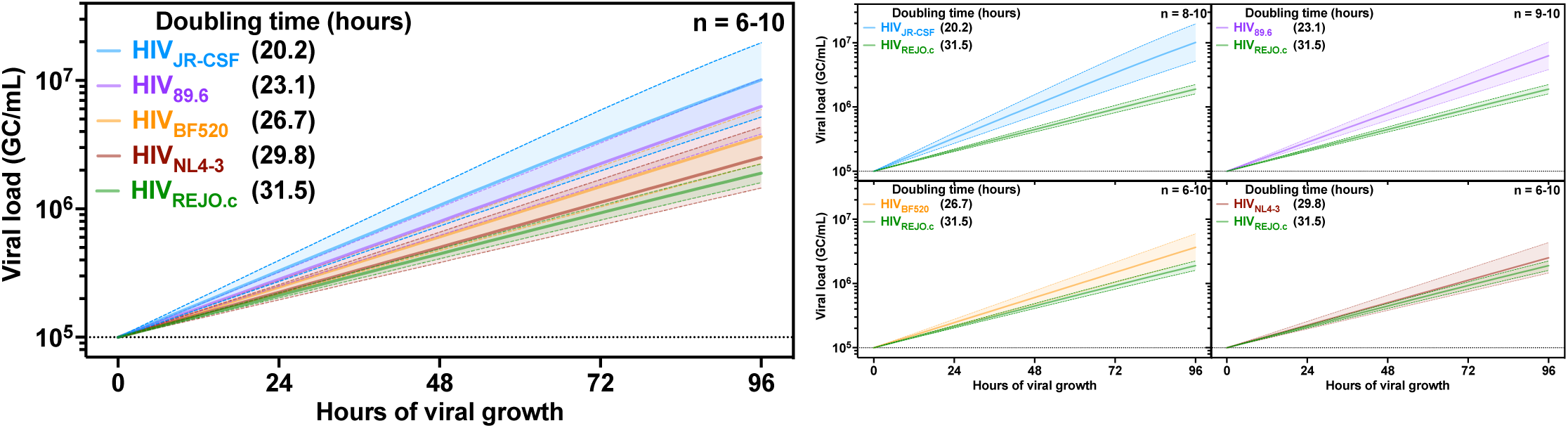
Different HIV strains exhibit different growth rates and carrying capacities. Comparison of HIV_REJO.c_, HIV_NL4-3_, HIV_JR-CSF_, HIV_89.6_, and HIV_BF520_ growth rates determined by QuickFit RT-qPCR. Viral growth (GC/mL) was calculated by normalizing the initial value on day 0 to 1×10^5^ GC/mL and subsequently interpolating the remaining values (filtered by inclusion and exclusion criteria) from days 1 to 4 based on the fit curve model. Overlaid is the mean growth rate and the 95% confidence interval. The Doubling time represents the number of hours it takes each isolate to double its viral load. The plots to the right show the pairwise comparison of each strain to HIV_REJO.c_. Dotted lines represent the limit of detection for each assay.

## DISCUSSION

Early efforts to determine HIV growth and fitness were based on kinetic data obtained in culture, using either individual viruses or or dual infection/growth competition experiments (37). Individual viral growth assays rely on measuring viral proteins, genetic material, physical particles, or reporter gene expression (38). These methods are usually time-consuming, expensive, and have different degrees of sensitivity between different viral isolates resulting in inconsistent growth rates (25–27,39). Dual infections or competitive growth assays employ a reference and a test strain to infect cells in the same culture, and the two strains are monitored via *env* gene heteroduplex tracking assays, allowing quantification of their growth and fitness (37,40). However, this fitness-measuring approach is time consuming and difficult to scale (41). More recent iterations of viral growth assays measure reverse transcriptase activity and perform linear regression analyses to determine growth kinetics, not only for HIV, but for lentiviruses in general (42,43). Other novel approaches measuring viral growth aim to streamline and high-throughput this process by performing parallel replication capacity assays followed by infection of reporter cell lines, such as TZM-bl, however these measurements are indirect (44). Dual infection cultures and allelic-specific PCRs can also be used to evaluate fitness differences between strains, yet these are difficult to scale efficiently (43,45,46).

Since a significant portion of viral growth measurements rely on ELISA-based approaches, we compared QuickFit to growth rates measured this way. ELISA assays have numerous sources of variability, including antibody concentrations, number of washes, sample dilution, and both the time and temperature of incubation for each step, leading to differences in results between plates (47,48). In our own studies, we found significant variation in the recognition of the HIV p24 capsid protein among different strains. Particularly, HIV_NL4-3_ exhibited the strongest signal, while HIV_BF520_ was only detectable at the highest inoculums, leading to wider measurement error. Protein alignment of the p24 capsid proteins of HIV_NL4-3_ and HIV_BF520_ results in pairwise identity of 91.3%, emphasizing that even relatively minor differences in antigens can have large impacts on the sensitivity of p24-based ELISAs (25). These issues can be minimized by incorporating standard curves to account for plate-to-plate variations, but may still contribute to the observed growth rates (49).

We developed QuickFit as a simpler and more robust method for the quantitation and analysis of viral growth and fitness, drawing from the advantages of previously reported approaches. The incorporation of a novel RNA extraction reagent enabled the use of high-throughput liquid handling in sample processing to minimize hands-on time (50–52). Measuring the growth rate of a single virus using QuickFit employs 80 distinct samples, representing approximately 20% of a 384-well plate. In contrast, p24 measurements obtained via ELISA require a full 96-well plate. As an RT-qPCR-based assay, QuickFit reliably measures even relatively small changes in fitness between different HIV strains (53,54). While we employed universal primers targeting highly conserved regions of the HIV *pol* gene, optimal sensitivity of these assays could be attained by precisely matching the sequence of the primers and probe to each strain (55). Not including the time needed to collect supernatants over multiple days, QuickFit can produce results in as little as 4 hours.

To reduce the intrinsic variability in viral growth that arises from *in vitro* culture, we analyzed QuickFit data on all 80 samples obtained across 16 wells for each individual virus to derive exclusion criteria that would minimize error. We excluded wells exhibiting decreasing viral load over time, as these were interpreted as failed cultures. We also excluded wells that reached the carrying capacity of the assay before the final time point, as well as any wells that failed to reach logarithmic growth, which has been shown to occur when the initial virus inoculum is too low (56,57). The need for these criteria to improve the reliability of QuickFit measurements is a consequence of using raw viral stocks, eliminating the need to titer each viral preparation.

Numerous studies have proposed mathematical models of HIV replication and growth determination *in vivo* and *in vitro* (37,56–59). The parameters considered in these models include the number of viral particles, the doubling time or growth rates, the carrying capacity, and the time span of the experiment. The use of a half-maximal equation via NLME modeling enables constraining some of these variables when required (60–62). In our case, when determining the growth rates for each individual well infected with the same HIV strain, we constrained the carrying capacity to a shared value (63,64). The same rubric was applied when we analyzed different HIV mutant strains derived from the same backbone, namely mutations in polymerase. Of note, analyzing these metrics for each Pol mutant strain independently resulted in somewhat different growth rates and carrying capacities as compared to those analyzed together. However, each metric was based on a limited number of wells that fell within the acceptable range of growth conditions, making them less reliable (64,65). When comparing growth rates of different HIV strains, more accurate values were obtained by allowing the NLME model to derive separate carrying capacities for each strain. The choice to constrain these variables should be made carefully when using QuickFit to ensure accurate estimates of growth rates.

Single-point mutations within the HIV *envelope* or *polymerase* gene can have profound effects on the fitness of HIV. This directly impacts the potential for HIV to escape from antibodies or antiretroviral drugs, as each mutation introduces a unique cost or benefit to viral growth (20). Previous reports suggest that the *pol* mutant M184V has an intermediate fitness relative to the WT and the M184I strains (32,66,67). In agreement with these findings, isoleucine at position M184 is observed more frequently in patients refractory to HAART at early time points, but valine quickly outcompetes it, becoming the dominant mutation in these patients over time (66,68,69). In our assays, M184I grew somewhat better than M184V in the absence of FTC; however, M184V grew substantially better than M184I in the presence of the drug. This shift in fitness in the presence or absence of drugs could account for the differences in relative fitness between mutants we observed via QuickFit compared to other reports (32).

Despite our efforts to optimize QuickFit, other sources of intrinsic error still remain, such as donor variability between PBMCs used for each assay (70). To address this, we evaluated the growth rates measured across three different donors and two HIV strains. Of note, the relative fitness cost within a given isolate (i.e., HIV_REJO.c_ WT and the Pol M184I mutant) remained similar, with only minor variations between donors. However, absolute growth rates of HIV_REJO.c_ and HIV_JR-CSF_ strains measured across different PBMCs were statistically different. This could be attributed to varying levels of co-receptor expression or host genetics, which could have impacted their susceptibility to infection (70). Despite this, by normalizing the growth rates of each donor, the relative difference between the HIV_REJO.c_ and the HIV_JR-CSF_ isolates was maintained. One potential solution to donor-to-donor variability would be to employ cell lines instead of primary PBMCs for HIV outgrowth assays (71,72). However, this presents additional caveats, as each cell line would exhibit different phenotypes relative to PBMCs, such as their surface CD4, CXCR4, and CCR5 expression levels, the internalization rates of these receptors upon HIV infection, and their intracellular dNTP availability (32,73).

Despite these caveats, QuickFit represents an improved platform for determining viral growth rates and evaluating fitness costs and could be further adapted to enhance its versatility. For instance, future iterations could utilize competitive growth assays with two or more strains grown in the same well, followed by multiplexed allele-specific RT-qPCR to provide individual measurements that are internally controlled. It could also be used to measure fitness of strains in outgrowth assays, using samples from animal models or people living with HIV to compare growth rates from diverse viral populations. Viral evolution studies and the selective pressure different drugs exert on the virus could also be evaluated with QuickFit. Finally, the use of genetically defined target cell lines could further reduce variability relative to PBMCs. In conclusion, QuickFit is a reproducible, quantitative, high-throughput method for the quantification of growth which could be useful for the evaluation of fitness costs imposed by antiviral drugs.

## Supplementary Materials

The following supporting information can be found online: Supplementary Tables 1-5; and Supplementary Figures 1-6.

## Author Contributions

Conceptualization, N.M.S.G, M.S., A.Z.L, and A.B.B.; methodology, N.M.S.G, M.S., A.Z.L., Y.C., E.C.L, A.M.J.; software, N.M.S.G, and A.Z.L; validation, N.M.S.G, M.S., A.Z.L., Y.C., E.C.L.; formal analysis, N.M.S.G, M.S., A.Z.L.; investigation, N.M.S.G, M.S., A.Z.L, and A.B.B.; resources, A.B.B.; data curation, N.M.S.G, M.S., A.Z.L, and A.B.B.; writing—original draft preparation, N.M.S.G, A.B.B.; writing—review and editing, N.M.S.G, M.S., A.Z.L., Y.C., A.M.J., and A.B.B.; visualization, N.M.S.G, M.S., and A.B.B.; supervision, A.B.B.; project administration, A.B.B; funding acquisition, A.B.B. All authors have read and agreed to the published version of the manuscript.

## Institutional Review Board Statement and Informed Consent Statement

Not applicable.

## Data Availability Statement

The raw data supporting the conclusions of this article will be made available by the authors upon request.

## Acknowledgments

The following reagents were obtained through the NIH HIV Reagent Program, Division of AIDS, NIAID, NIH: HIV-1, Strain NL4-3 Infectious Molecular Clone (pNL4-3), ARP-114, contributed by Dr. M. Martin; HIV-1, Strain JR-CSF Infectious Molecular Clone (pYK-JRCSF), ARP-2708, contributed by Dr. Irvin S. Y. Chen and Dr. Yoshio Koyanagi; HIV-1, 89.6 Infectious Molecular Clone (p89.6), ARP-3552, contributed by Dr. Ronald G. Collman; ARP-11746 (pREJO.c/2864), contributed by Dr. John Kappes and Dr. Christina Ochsenbauer; TZM-bl Cells, ARP-8129, contributed by Dr. John C. Kappes, Dr. Xiaoyun Wu and Tranzyme Inc. We thank J. Bloom for providing the HIV_BF520_ infectious molecular clone plasmids. Quantitation of p24 concentration was performed with reagents provided by the AIDS and Cancer Virus Program, Leidos Biomedical Research, Inc., Frederick National Laboratory for Cancer Research, supported with federal funds from the National Cancer Institute, National Institutes of Health, under contract HHSN261200800001E

## MATERIALS AND METHODS

### Isolation and Activation of CD4^+^ T cells

Primary human PBMCs were isolated from human blood samples by Ficoll density gradient centrifugation, as reported previously (74). Briefly, blood was diluted 1:2 in 1x DPBS (Corning, #21031CV) and gently layered over Ficoll buffer (Cytiva, #17144002) in a 50 ml conical tube. Diluted blood was then spun at 400 x g for 40 min. The resulting monolayer of cells was isolated and washed using RPMI1640 (Stemcell, #36750) supplemented with 10% FBS (VWR, #97068-085), Pen/Strep (Thermo, #15140122), L-Glutamax (Life Technologies, #25030081), MEM Non-essential amino acids (Corning, #25-025-CI), HEPES (Thermo, #MT25060CI), and Sodium Pyruvate (Corning, 25-000-CI) (isolation media). PBMCs were counted and frozen down in 90% FBS and 10% DMSO at a density of 20×10^6^ cells per tube. Then, CD4^+^ T cells were isolated from frozen PBMC tubes using the EasySep™ Human CD4^+^ T Cell Isolation Kit (Stem Cell Technologies, #17952) following the manufacturer’s protocol. Isolated naive CD4^+^ T cells were resuspended in isolation media supplemented with recombinant IL-2 (R&D Systems, #202-IL-050) at 10 ng/mL (growth media) and 4 µg/mL of anti-CD28 antibody (Biolegend, #302934). Cells were plated in non-TC treated 24-well plates previously coated with 2 µg/mL of anti-CD3 antibody (Biolegend, #317304) overnight at 4°C. Cells were incubated at 37°C and 5% CO_2_ for 2 to 4 days until proliferation was visible. Cells were then washed with isolation media and pooled with growth media in a T75 flask for another 2-4 days before using in downstream assays. The purification and activation efficiency were routinely assessed by flow cytometry. All the experiments presented here were performed using PBMCs from the same donor in order to reduce variability unless stated otherwise.

### Infection of Primary Human CD4^+^ T cells with HIV-1 Infectious Molecular Clones

HIV-1 viral stocks were obtained by transfecting 293T cells (ATCC, #CRL-3216) with full-length replication competent infectious molecular clone (IMC) plasmids, as described previously (BEI Resources, #ARP114, #ARP2708, ARP#3552, #ARP11746, #ARP13402) (75).

All IMC plasmids (both WT and mutants) were routinely full-length sequenced to confirm their accuracy. The TCID_50_ and infectivity of viral stocks were routinely assessed on TZM-bl cells (BEI resources, #ARP8129) prior to any assay, as previously described (76–80). HIV-1 viral stocks were then diluted in duplicate in growth media in 96-well flat bottom plates. Activated CD4^+^ T cells were added to the plate at a density of 1×10^5^ cells/well. Plates were then spinoculated at 400 x g for 1 h and incubated overnight at 37°C and 5% CO_2_. After 12h, the cells were transferred to 96-well round bottom plates and washed 3 times with isolation media to remove unbound viral particles by spinning them down at 500 x g for 5 min and removing the media. After washing, the cells were resuspended in 200 µL of growth media and incubated at 37°C and 5% CO_2_ for six days. If samples were treated with emtricitabine (Emtriva^TM^ FTC Gilead, #61958-0601), growth media was supplemented with 70 ng/mL in all steps. As a vehicle control, the same volume of DMSO was added to the growth media of cells that were not treated with FTC.

### Supernatant collection and RNA extraction

Viral supernatants were collected starting 48 h after the cells were transferred to round bottom 96-well plates and until the end of the assay. A total of 24 µL was collected daily by a TECAN Fluent 780 Liquid Handler and fresh media with the same composition as the one collected was added back to the plates to recover the original volume. Upon collection, supernatants were immediately diluted 4:1 into QuickExtract DNA Extraction Solution (Biosearch Technologies, #QE09050), homogenized, and incubated in a thermocycler at 95°C for 5 min and 4°C for 5 min (50–52). Alternatively, supernatants were immediately frozen so protein lysis could be performed later. Extracted viral RNA was then stored at -80°C until RT-qPCR were performed. QuickExtract DNA Extraction Solution freeze-thaw cycles were limited to no more than 3 to prevent decreased effectiveness.

### p24 ELISA quantification

P24 capsid concentration in viral supernatants was quantified with the Leidos Biomedical Research HIV-1 p24CA Antigen Capture Assay Kit, following the manufacturer’s instructions. Briefly, 96-well ELISA plates (Thermo #446612) were coated with a 1:400 dilution of the capture antibody (Lot#PP292-3) in DPBS and incubated overnight at 4°C. Plates were then blocked for one hour and washed six times with wash buffer. During this incubation, frozen samples were thawed, and protein lysis was performed by adding 1µL of 10% Triton X-100 (Thermo #BP151-500) in dH_2_O to 19µL of sample and then incubating this for 1 h at 37°C. Samples were diluted in sample diluent, added to the plates, and incubated for 2 h at 37°C. All samples were run in duplicate. Independent standard curves diluted in sample diluent were held for each plate (Lot#SP968T). The plates were washed as before, and then a 1:300 dilution of the primary antibody (Lot#SP2143A) in the corresponding diluent was added to the plates. Plates were incubated for 1 h at 37°C, washed six times again, and a 1:25,000 dilution of the secondary antibody (Goat anti-Rabbit IgG H+L Chain HRP Conjugated - Bethyl #A120-201P) in the corresponding diluent was added to the plates. The optimal secondary antibody concentration was previously determined by testing several dilutions of this reagent with dilutions of the standard curve. After incubating for 1 h at 37°C, plates were washed one last time as before and one time with DPBS. For the readouts, 100 µL of TMB substrate (VWR #95059-156) was added to the wells and incubated for 30 min at room temperature. The reactions were stopped by adding 100 µL of 1N HCL, and the OD_450_ and OD_650_ values were read within five minutes in an Emax Laboratory Precision Microplate Reader. The OD_650_ was subtracted from the OD_450_, and p24 concentrations were determined by interpolating these values into each independent standard curve using a four-parameter equation.

### Quantification of viral gene copies by RT-qPCR

Extracted viral RNA was thawed, and 5 µL were used in a 10 µL RT-qPCR reaction with the qScript XLT one-step RT-qPCR Tough Mix, low ROX kit (Quanta Biosciences, #95134-500), a TaqMan probe (56-FAM/CCCACCAAC/ZEN/AGGCGGCCTTAACTG/3IABkFQ) and primers designed to target the *pol* gene of HIV_REJO.c_, (5’-CAATGGCCCCAATTTCATCA and 3’-GAATGCCGAATTCCTGCTTGA), HIV_NL4-3_/HIV_JR-CSF_/HIV_89.6_(5’-CAATGGCAGCAATTTCACCA and 3’-GAATGCCAAATTCCTGCTTGA) and HIV_BF520_ (5’-CAATGGCAGCAATTTCACCA and 3’-GAATCCCAAATTCCTGTTGGA) (Integrate DNA technologies). A different TaqMan probe to detect HIV_BF520_ was used (56-FAM/CCCACCAAC/ZEN/AGGCTGCTTTAACTG/ZEN/3IABkFQ/). Samples were run on a QuantStudio 12K Flex (Applied Biosystems). The following cycling conditions were used: 50°C for 10 min, 95°C for 3 min, followed by 55 cycles of 95°C for 3 s and 60°C for 30 s. Viral loads (Genome copies (GC)/mL) were determined by interpolating the CT values into a standard curve generated with RNA extracted from a previously titered serially-diluted viral stock. The range of the assay was 1×10^4^ to 1×10^9^ GC/mL.

### Growth rate quantitation and growth curve modeling

To quantify growth rates and model growth curves, the viral growth indicator data (GC/mL or p24 pg/mL) was either filtered or not with the following set of rules: For the p24 ELISA data, any individual wells not reaching a value above the limit of detection on day 2, those reaching the upper limit of detection at any time, and those where p24 concentration decreased up to a 10% relative to the previous quantitation were excluded; for the RT-qPCR data, any individual wells reaching the lower or upper limit of detection at any time point, those where viral loads decreased up to 10% relative to the previous quantitation, and those where the initial viral load was more than 0.25 times or less than 0.025 the highest quantified value for that well were excluded. Viral growth indicator values were corrected to account for the corresponding dilution factor for each day after sample collection.

After filtering, viral growth indicator data was analyzed using the non-linear mixed-effect (NLME) modeling software MonolixSuite 2021R (Lixoft SAS, Antony, France) (60,61). The following equation was used to estimate growth rates (r) and carrying capacities (K) population parameters using a continuous observation model type.

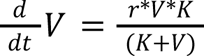

**Equation 1**: Half-maximal formula used to quantify growth rates.

With *V* being the quantified viral growth indicator data and *t* being the time. For the estimation of the r values, random effects were allowed for each individual model. The estimation of the individual K values was constrained so that they would remain constant for each HIV isolate unless stated otherwise. The output viral growth indicator data obtained was used to model growth curves as it was or by normalizing it to the initial values obtained for each individual sample. Data was then plotted using GraphPad Prism v10.2.2 (LLC).

## SUPPLEMENTARY FIGURE LEGENDS

**Supplementary Figure 1.**
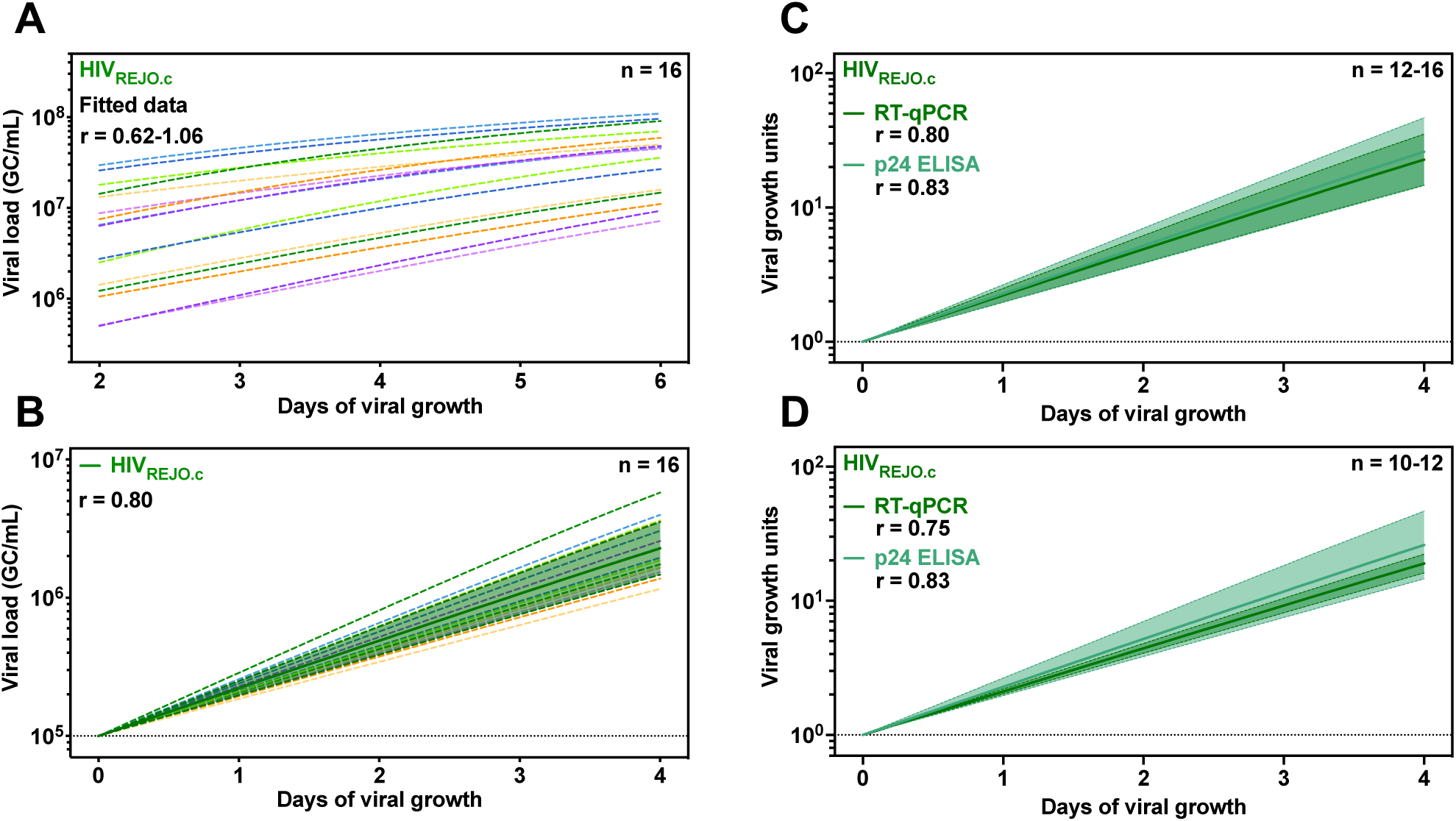
Analysis and modeling of HIV_REJO.c_ *in vitro* replication without data filtering and comparison of ELISA and QuickFit growth curves. (**A**) Viral loads (GC/mL) were determined for HIV_REJO.c_ by QuickFit RT-qPCR and all the data from each individual well were used to determine growth rates and carrying capacities by running an NMLE modeling with a half-maximal equation in the MonolixSuite software. A total of 16 samples at 5 different time points were evaluated, and matching light-dark colors represent duplicates. (**B**) Viral growth (GC/mL) was calculated by normalizing the initial value on day 0 to 1×10^5^ GC/mL and subsequently interpolating the remaining values from days 1 to 4 based on the fit curve model. Overlaid is the mean growth rate and the 95% confidence interval. (**C**) Viral growth (p24 concentration or GC/mL) of inclusion and exclusion criteria filtered data was calculated by normalizing the initial value on day 0 to either 1.00 pg/mL or 1.00 GC/mL and subsequently interpolating the remaining values from days 1 to 4 based on the fit curve model. Overlaid is the mean growth rate and the 95% confidence interval. The r value represents the average growth rate determined for all included samples by each assay.

**Supplementary Figure 2.**
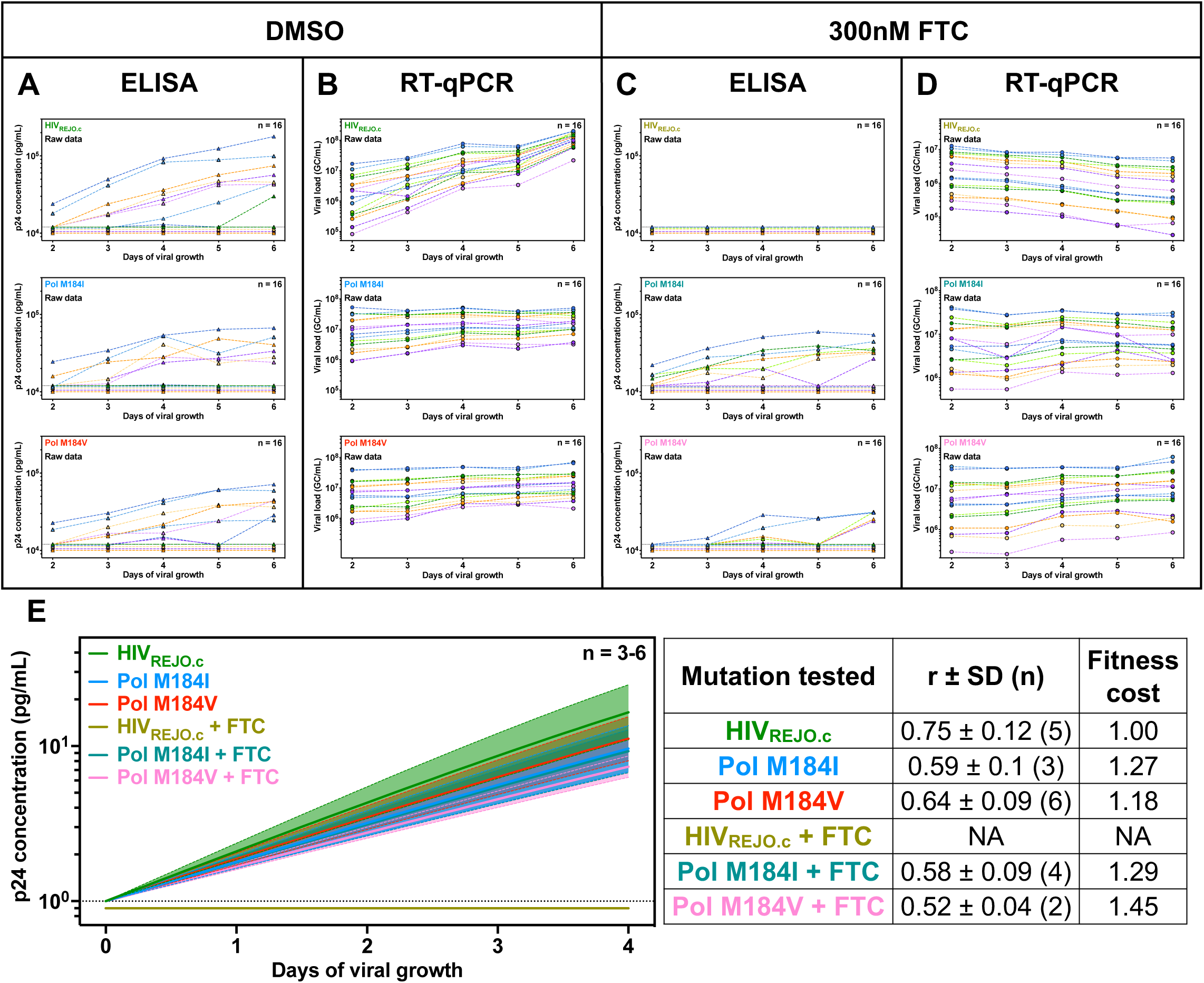
Determination of HIV_REJO.c_ WT and Pol mutants raw p24 concentration and viral loads and evaluation of growth rates by p24 ELISA. Single-point mutations for the pol gene were introduced in the HIV_REJO.c_ IMC backbone and p24 ELISA and RT-qPCR were performed with samples obtained from the QuickFit pipeline experimental setup. These strains were also grown in the presence of 300nM emtricitabine (FTC) or DMSO as a vehicle control. Determined raw p24 concentrations (pg/mL) for the different strains grown in (**A**) DMSO- or (**C**) FTC - containing media and raw viral loads (GC/mL) for the different strains grown in (**B**) DMSO- or (**D**) FTC - containing media are shown. A total of 16 samples at 5 different time points were evaluated. The limits of detection (LOD) shown correspond to the standard curve working range after sample dilution correction. (**E**) Viral growth (p24 concentration) was calculated by normalizing the initial value of each sample (after inclusion and exclusion criteria filtering) on day 0 to 1.00 pg/mL and subsequently interpolating the remaining values from days 1 to 4 based on the fit curve model. Overlaid is the mean growth rate and the 95% confidence interval. The table to the right shows the r value (growth rates) ± the standard deviation for each strain tested for both conditions. The sample size (n) references the number of wells included in the analysis after filtering with inclusion and exclusion criteria. The fitness cost represents the doubling rate of each strain relative to the WT HIV_REJO.c_ without FTC. **NA**: No growth was detected.

**Supplementary Figure 3.**
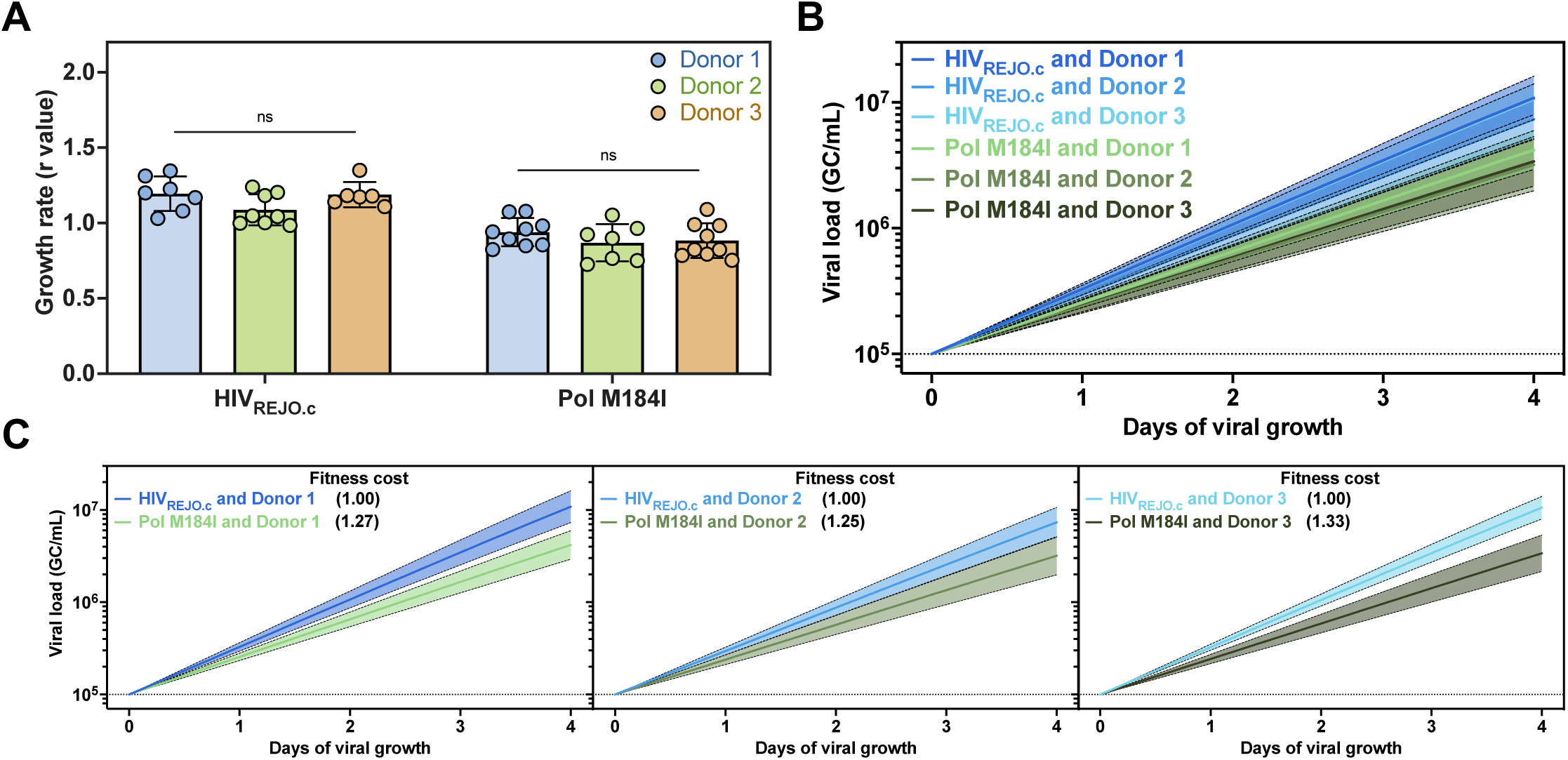
Changes in fitness cost for the same isolate are retained despite donor variability. The growth rates for the HIV_REJO.c_ WT and Pol M184I mutant strains were evaluated using different PBMC donors and constraining the simulations to a shared K value. (**A**) Comparison of r values determined for both strains for each donor (ns: non-significant; Two-way ANOVA with Šídák *post hoc* analysis). (**B**) Normalized growth curves were generated for both strains and for each donor independently. (**C**) Pairwise comparison of the normalized growth curves generated for both strains and for each donor. The fitness cost represents the doubling rate of each strain relative to the WT HIV_REJO.c_.

**Supplementary Figure 4.**
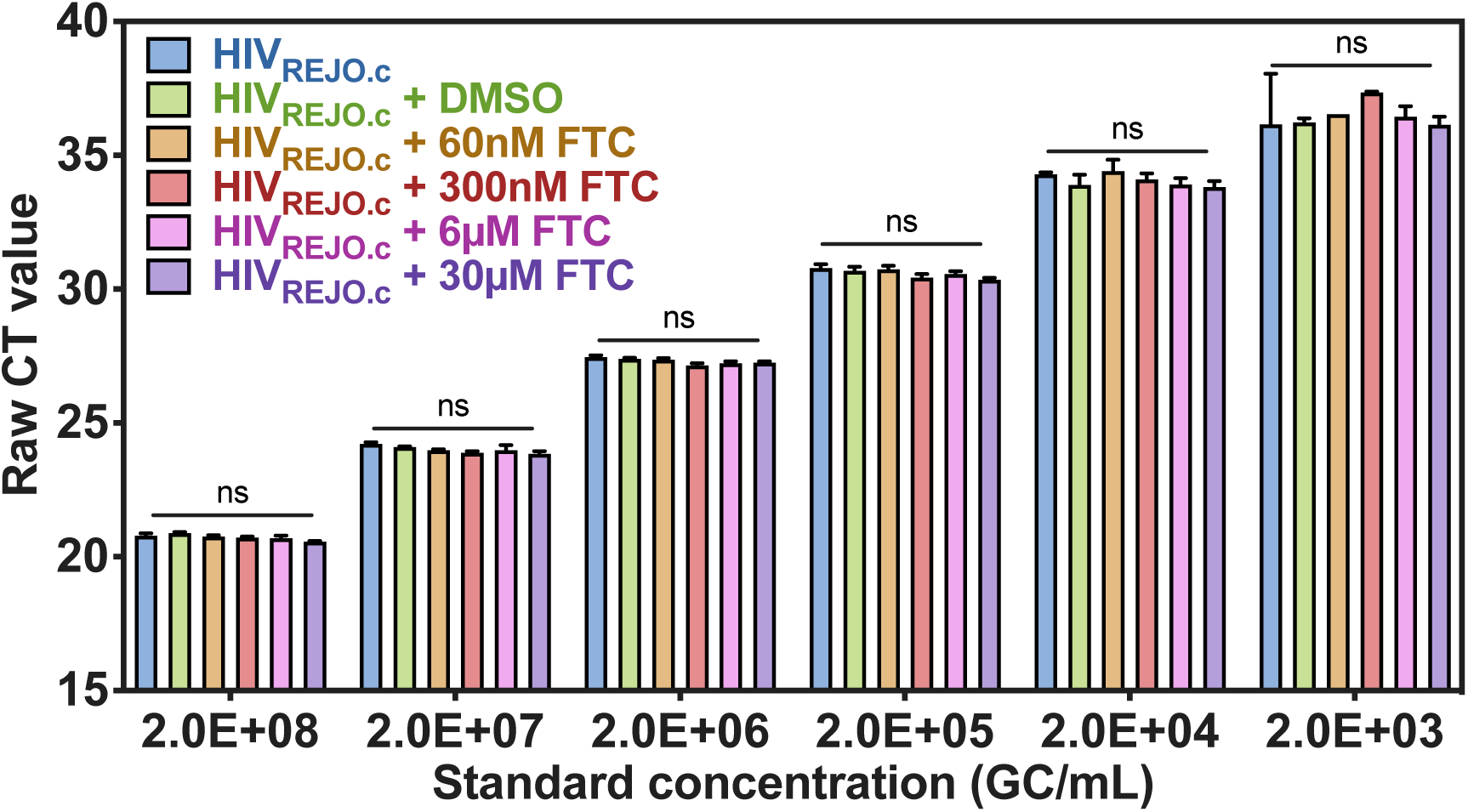
Emtricitabine does not impact the RT-qPCR quantification. Serial dilutions of a previously tittered stock of HIV_REJO.c_ were quantified by RT-qPCR in the presence of increasing concentrations of emtricitabine (FTC, an RT inhibitor) ranging from 35 ng/mL (60nM) to 700 ng/mL (30µM). Raw CT values measured are shown. The highest FTC concentration tested here is 10 times higher than the one tested in the experiments shown in **Figure 3**. No statistically significant differences were found for any of the different FTC concentrations at any specific standard concentrations (Two-way ANOVA with Šídák *post hoc* analysis).

**Supplementary Figure 5.**
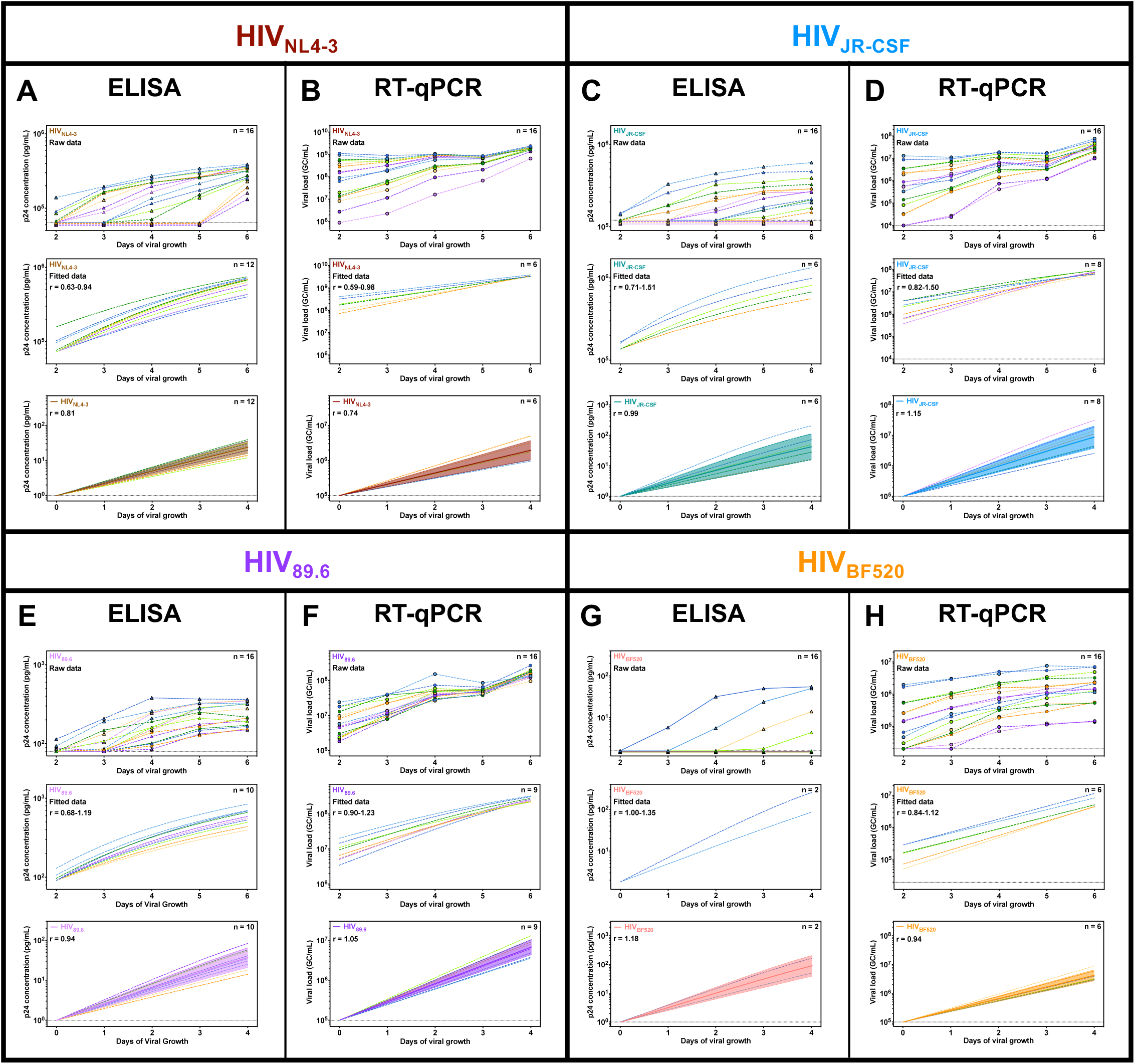
Analysis and modeling of in vitro replication growth curves for four different HIV-1 strains as measured by p24 ELISA and QuickFit RT-qPCR. Both p24 ELISA (**left**) and QuickFit RT-qPCR (**right**) were performed for the four different HIV-1 strains (**A,B**) HIV_NL4-3_, (**C,D**) HIV_JR-CSF_, (**E,F**) HIV_89.6_, and (**G,H**) HIV_BF520_. (**A,C,E,G, top**) Determined raw p24 concentrations (pg/mL) and (**B,D,F,H, top**) viral loads (GC/mL) are shown. A total of 16 samples at 5 different time points were evaluated. The limits of detection (LOD) shown correspond to the standard curve working range after sample dilution correction. (**Middle**) Samples were filtered using inclusion and exclusion criteria, and the remaining data were used to determine growth rates and carrying capacities by running an NMLE modeling with a half-maximal equation in the MonolixSuite software. (Bottom) Normalized viral growth (p24 concentration or GC/mL) was calculated by normalizing the initial value on day 0 to 1.00 pg/mL or 1×10^5^ GC/mL and subsequently interpolating the remaining values from days 1 to 4 based on the fit curve model. Overlaid is the mean growth rate and the 95% confidence interval. The r value represents the average growth rate determined for all included samples by each assay.

**Supplementary Figure 6.**
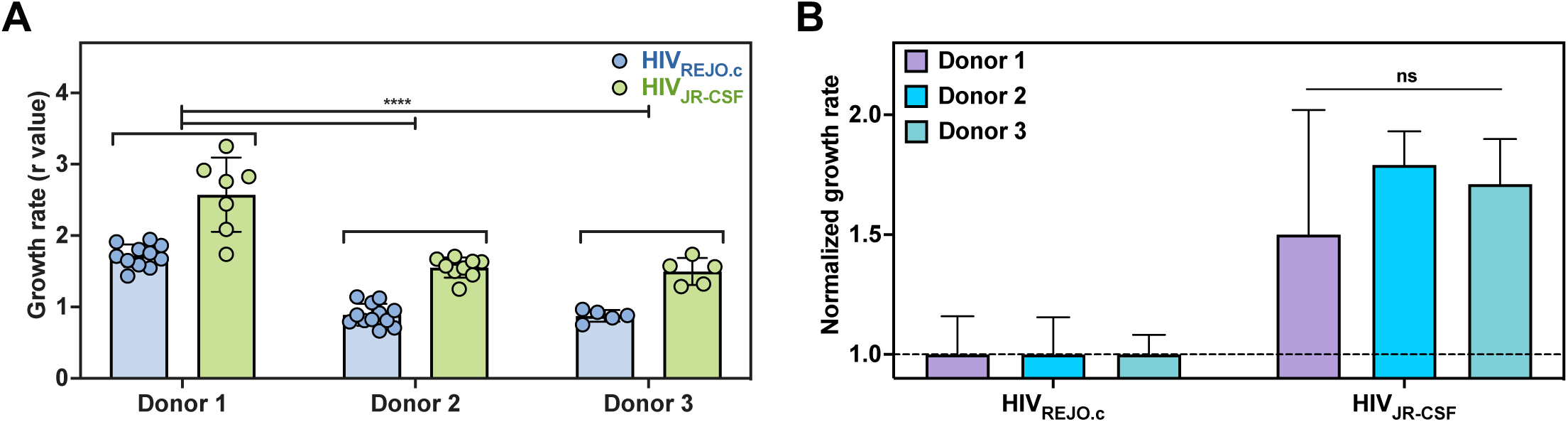
Growth rates for different isolates may vary between donors. The growth rates for HIV_REJO.c_ and HIV_JR-CSF_ were determined using PBMCs from three different donors. A single simulation, including all strains and all donors, was performed without constraining the carrying capacity to a shared value. (**A**) Comparison of growth rates between each donor (**\*\*\*\*:** p<0.001; Two-way ANOVA with Šídák *post hoc* analysis). (**B**) The growth rate of HIV_JR-CSF_ for each donor was normalized to that of the corresponding HIV_REJO._c and then compared between each other to determine whether the doubling rate ratio was maintained across donors (ns: non-significant; Two-way ANOVA with Šídák *post hoc* analysis).

