## Supplementary Tables 1-5 for "QuickFit: A high-throughput RT-qPCR-based assay to quantify viral growth and fitness *in vitro*"

**Supplementary Table 1. Raw p24 concentration and viral loads.**

| p24 ELISA Raw pg/mL | | | | | | | | | | |
|---|---|---|---|---|---|---|---|---|---|---|
| HIV REJO.c | Replicate 1 | | | | | Replicate 2 | | | | |
| Dilution | Day 2 | Day 3 | Day 4 | Day 5 | Day 6 | Day 2 | Day 3 | Day 4 | Day 5 | Day 6 |
| 1:2 | 2.48E+04 | 5.78E+04 | 1.43E+05 | 2.25E+05 | 2.92E+05 | 2.42E+04 | 6.10E+04 | 1.25E+05 | 2.22E+05 | 2.97E+05 |
| 1:4 | 1.60E+04 | 3.86E+04 | 9.07E+04 | 1.60E+05 | 2.20E+05 | 1.60E+04 | 3.73E+04 | 9.57E+04 | 1.59E+05 | 2.11E+05 |
| 1:8 | 1.60E+04 | 2.72E+04 | 7.08E+04 | 1.24E+05 | 1.88E+05 | 1.60E+04 | 2.56E+04 | 6.96E+04 | 1.38E+05 | 1.94E+05 |
| 1:16 | 1.60E+04 | 1.83E+04 | 5.31E+04 | 1.01E+05 | 1.36E+05 | 1.60E+04 | 1.84E+04 | 5.71E+04 | 1.24E+05 | 1.87E+05 |
| 1:32 | 1.60E+04 | 1.60E+04 | 3.30E+04 | 6.70E+04 | 9.91E+04 | 1.60E+04 | 1.60E+04 | 3.49E+04 | 7.85E+04 | 1.23E+05 |
| 1:64 | 1.60E+04 | 1.60E+04 | 2.96E+04 | 6.57E+04 | 1.06E+05 | 1.60E+04 | 1.60E+04 | 2.17E+04 | 5.64E+04 | 1.02E+05 |
| 1:128 | 1.60E+04 | 1.60E+04 | 1.60E+04 | 4.28E+04 | 7.35E+04 | 1.60E+04 | 1.60E+04 | 1.60E+04 | 3.99E+04 | 6.69E+04 |
| 1:256 | 1.60E+04 | 1.60E+04 | 1.60E+04 | 1.60E+04 | 2.23E+04 | 1.60E+04 | 1.60E+04 | 1.60E+04 | 2.98E+04 | 5.79E+04 |
| RT-qPCR Raw GC/mL | | | | | | | | | | |
| HIV REJO.c | Replicate 1 | | | | | Replicate 2 | | | | |
| Dilution | Day 2 | Day 3 | Day 4 | Day 5 | Day 6 | Day 2 | Day 3 | Day 4 | Day 5 | Day 6 |
| 1:2 | 2.28E+07 | 3.41E+07 | 3.51E+07 | 5.14E+07 | 5.58E+07 | 2.61E+07 | 4.27E+07 | 5.14E+07 | 4.41E+07 | 7.07E+07 |
| 1:4 | 1.26E+07 | 2.49E+07 | 4.14E+07 | 4.01E+07 | 4.44E+07 | 1.58E+07 | 2.01E+07 | 2.98E+07 | 2.51E+07 | 4.06E+07 |
| 1:8 | 6.62E+06 | 1.43E+07 | 2.35E+07 | 2.18E+07 | 2.93E+07 | 1.16E+07 | 1.58E+07 | 1.80E+07 | 1.91E+07 | 2.38E+07 |
| 1:16 | 5.60E+06 | 1.15E+07 | 1.77E+07 | 1.47E+07 | 2.56E+07 | 7.70E+06 | 1.12E+07 | 1.71E+07 | 1.73E+07 | 2.23E+07 |
| 1:32 | 2.42E+06 | 3.25E+06 | 8.51E+06 | 7.75E+06 | 1.51E+07 | 5.72E+06 | 7.04E+06 | 1.60E+07 | 1.42E+07 | 2.47E+07 |
| 1:64 | 1.08E+06 | 1.96E+06 | 4.01E+06 | 4.61E+06 | 7.08E+06 | 2.21E+06 | 4.45E+06 | 1.07E+07 | 8.07E+06 | 1.53E+07 |
| 1:128 | 9.31E+05 | 1.07E+06 | 2.85E+06 | 3.91E+06 | 5.41E+06 | 1.26E+06 | 1.24E+06 | 4.19E+06 | 4.97E+06 | 9.04E+06 |
| 1:256 | 4.42E+05 | 6.66E+05 | 2.37E+06 | 2.52E+06 | 3.31E+06 | 4.50E+05 | 6.39E+05 | 1.89E+06 | 1.79E+06 | 3.46E+06 |

**†Greyed-out rows represent sample sets excluded from downstream analyses according to inclusion and exclusion criteria.**

**Supplementary Table 2. Individual and average growth rates and carrying capacities determined for $HIV_{REJO.c}$ by p24 ELISA and RT-qPCR with and without data filtering.**

| $HIV_{REJO.c}$ | p24 ELISA | | Unfiltered RT-qPCR Data | | Filtered RT-qPCR Data | |
|---|---|---|---|---|---|---|
| **K (pg/mL or GC/mL)*** | 5.96E+05 | | 3.16E+07 | | 3.85E+07 | |
| **Dilution** | **Replicate 1** | **Replicate 2** | **Replicate 1** | **Replicate 2** | **Replicate 1** | **Replicate 2** |
| **1:2** | 0.99 | 0.99 | 0.88 | 0.95 | | |
| **1:4** | 0.99 | 0.99 | 1.06 | 0.75 | | |
| **1:8** | 0.88 | 0.89 | 0.92 | 0.62 | 0.81 | |
| **1:16** | 0.77 | 0.85 | 0.83 | 0.69 | 0.76 | |
| **1:32** | 0.64 | 0.69 | 0.76 | 0.80 | 0.74 | 0.75 |
| **1:64** | 0.64 | 0.61 | 0.73 | 0.92 | 0.72 | 0.82 |
| **1:128** | | | 0.67 | 0.72 | 0.69 | 0.71 |
| **1:256** | | | 0.80 | 0.71 | 0.77 | 0.71 |
| **Average r ± SD**** | **0.83 ± 0.14** | | **0.80 ± 0.12** | | **0.75 ± 0.04** | |

***Carrying capacity values (K) correspond to the average highest value Viral Growth indicator (either p24 pg/mL or GC/mL) that could be sustained in a steady state.**
****The r value represents the replication rate measured for each data set.**
**†Greyed-out rows represent sample sets excluded from downstream analyses according to inclusion and exclusion criteria.**

**Supplementary Table 3. Individual and average growth rates and carrying capacities determined for HIV$_{REJO.c}$ WT and Pol mutants by p24 ELISA and RT-qPCR after data filtering.**

| HIV$_{REJO.c}$ | ELISA | | Filtered qPCR Data | | Pol M184I | ELISA | | Filtered qPCR Data | | Pol M184V | ELISA | | Filtered qPCR Data | |
|---|---|---|---|---|---|---|---|---|---|---|---|---|---|---|
| K (pg/mL or GC/mL)* | 8.00E+04 | | 1.70E+07 | | K (pg/mL or GC/mL)* | 8.00E+04 | | 1.70E+07 | | K (pg/mL or GC/mL)* | 8.00E+04 | | 1.70E+07 | |
| Dilution | Replicate 1 | Replicate 2 | Replicate 1 | Replicate 2 | Dilution | Replicate 1 | Replicate 2 | Replicate 1 | Replicate 2 | Dilution | Replicate 1 | Replicate 2 | Replicate 1 | Replicate 2 |
| 1:2 | 0.93 | 0.77 | | | 1:2 | 0.71 | | | | 1:2 | 0.71 | 0.73 | | |
| 1:4 | 0.77 | 0.70 | | | 1:4 | 0.55 | 0.53 | | | 1:4 | 0.63 | 0.68 | | |
| 1:8 | 0.59 | | | | 1:8 | | | | | 1:8 | | 0.58 | | 0.59 |
| 1:16 | | | 1.01 | 0.79 | 1:16 | | | | | 1:16 | | 0.48 | 0.56 | 0.47 |
| 1:32 | | | 0.88 | 0.86 | 1:32 | | | 0.62 | 0.64 | 1:32 | | | 0.59 | 0.48 |
| 1:64 | | | 0.81 | 0.92 | 1:64 | | | 0.64 | 0.60 | 1:64 | | | 0.57 | 0.61 |
| 1:128 | | | 0.73 | 0.81 | 1:128 | | | 0.66 | 0.71 | 1:128 | | | 0.55 | 0.51 |
| 1:256 | | | | | 1:256 | | | | 0.70 | 1:256 | | | | |
| Average r ± SD** | 0.75 ± 0.12 | | 0.85 ± 0.09 | | Average r ± SD | 0.59 ± 0.1 | | 0.65 ± 0.04 | | Average r ± SD | 0.64 ± 0.09 | | 0.55 ± 0.05 | |
| HIV$_{REJO.c}$ + FTC | ELISA | | Filtered qPCR Data | | Pol M184I + FTC | ELISA | | Filtered qPCR Data | | Pol M184V + FTC | ELISA | | Filtered qPCR Data | |
| K (pg/mL or GC/mL)* | 8.00E+04 | | 1.70E+07 | | K (pg/mL or GC/mL)* | 8.00E+04 | | 1.70E+07 | | K (pg/mL or GC/mL)* | 8.00E+04 | | 1.70E+07 | |
| Dilution | Replicate 1 | Replicate 2 | Replicate 1 | Replicate 2 | Dilution | Replicate 1 | Replicate 2 | Replicate 1 | Replicate 2 | Dilution | Replicate 1 | Replicate 2 | Replicate 1 | Replicate 2 |
| 1:2 | | | | | 1:2 | 0.69 | 0.58 | | | 1:2 | 0.55 | 0.49 | | |
| 1:4 | | | | | 1:4 | 0.58 | 0.47 | | | 1:4 | | | | |
| 1:8 | | | | | 1:8 | | | | | 1:8 | | | | 0.51 |
| 1:16 | | | | | 1:16 | | | | | 1:16 | | | 0.54 | 0.51 |
| 1:32 | | | | | 1:32 | | | 0.35 | 0.38 | 1:32 | | | 0.43 | 0.45 |
| 1:64 | | | | | 1:64 | | | 0.42 | | 1:64 | | | 0.49 | 0.50 |
| 1:128 | | | | | 1:128 | | | | | 1:128 | | | | |
| 1:256 | | | | | 1:256 | | | | | 1:256 | | | | |
| Average r ± SD | NA | | NA | | Average r ± SD | 0.58 ± 0.09 | | 0.38 ± 0.03 | | Average r ± SD | 0.52 ± 0.04 | | 0.49 ± 0.04 | |

**[*]Carrying capacity values (K) correspond to the average highest value Viral Growth indicator (either p24 pg/mL or GC/mL) that could be sustained in a steady state.**
**[**]The r value represents the replication rate measured for each data set.**
**[†]Greyed-out rows represent sample sets excluded from downstream analyses according to inclusion and exclusion criteria.**

**Supplementary Table 4. Individual and average growth rates and carrying capacities determined for different HIV strains by p24 ELISA and RT-qPCR after data filtering.**

| $HIV_{NL4-3}$ | ELISA | | Filtered qPCR Data | | $HIV_{JR-CSF}$ | ELISA | | Filtered qPCR Data | |
|---|---|---|---|---|---|---|---|---|---|
| **K (pg/mL or GC/mL)*** | 4.09E+05 | | 4.53E+11 | | **K (pg/mL or GC/mL)*** | 2.97E+05 | | 1.14E+08 | |
| **Dilution** | **Replicate 1** | **Replicate 2** | **Replicate 1** | **Replicate 2** | **Dilution** | **Replicate 1** | **Replicate 2** | **Replicate 1** | **Replicate 2** |
| **1:2** | 0.87 | 0.75 | | | **1:2** | | | | |
| **1:4** | 0.92 | 0.87 | | | **1:4** | | | 0.82 | 0.96 |
| **1:8** | 0.94 | 0.92 | 0.59 | 0.56 | **1:8** | 1.13 | 1.51 | 0.94 | 1.13 |
| **1:16** | 0.88 | 0.83 | 0.74 | 0.72 | **1:16** | 0.86 | 1.02 | 1.20 | 1.27 |
| **1:32** | 0.67 | 0.76 | 0.98 | 0.87 | **1:32** | 0.71 | 0.71 | 1.35 | 1.50 |
| **1:64** | 0.66 | 0.63 | | | **1:64** | | | | |
| **1:128** | | | | | **1:128** | | | | |
| **1:256** | | | | | **1:256** | | | | |
| **Average r ± SD** | **0.81 ± 0.11** | | **0.74 ± 0.16** | | **Average r ± SD** | **0.99 ± 0.3** | | **1.15 ± 0.23** | |
| $HIV_{89.6}$ | **ELISA** | | **Filtered qPCR Data** | | $HIV_{BF520}$ | **ELISA** | | **Filtered qPCR Data** | |
| **K (pg/mL or GC/mL)*** | 2.46E+05 | | 3.52E+08 | | **K (pg/mL or GC/mL)*** | 5.40E+05 | | 7.00E+09 | |
| **Dilution** | **Replicate 1** | **Replicate 2** | **Replicate 1** | **Replicate 2** | **Dilution** | **Replicate 1** | **Replicate 2** | **Replicate 1** | **Replicate 2** |
| **1:2** | 0.88 | 1.19 | | | **1:2** | 1.35 | 1.00 | | |
| **1:4** | 0.74 | 1.07 | 0.97 | 0.91 | **1:4** | | | | |
| **1:8** | 0.68 | 1.11 | 1.04 | 0.90 | **1:8** | | | 0.92 | 0.84 |
| **1:16** | | 1.05 | 1.16 | 1.10 | **1:16** | | | 0.85 | 0.87 |
| **1:32** | | 0.97 | 1.13 | 1.04 | **1:32** | | | 1.02 | 1.12 |
| **1:64** | | 0.83 | 1.23 | | **1:64** | | | | |
| **1:128** | | 0.92 | | | **1:128** | | | | |
| **1:256** | | | | | **1:256** | | | | |
| **Average r ± SD** | **0.94 ± 0.16** | | **1.05 ± 0.11** | | **Average r ± SD** | **1.18 ± 0.25** | | **0.94 ± 0.11** | |

**[*]Carrying capacity values (K) correspond to the average highest value Viral Growth indicator (either p24 pg/mL or GC/mL) that could be sustained in a steady state.**
**[**]The r value represents the replication rate measured for each data set.**
**[†]Greyed-out rows represent sample sets excluded from downstream analyses according to inclusion and exclusion criteria.**

**Supplementary Table 5. Individual and average growth rates and carrying capacities determined for different HIV strains by RT-qPCR running a single simulation.**

| HIV Strain | $HIV_{REJO.c}$ | | $HIV_{NL4-3}$ | | $HIV_{JR-CSF}$ | | $HIV_{89.6}$ | | $HIV_{BF520}$ | |
|---|---|---|---|---|---|---|---|---|---|---|
| K (GC/mL)* | 2.34E+07 | | 9.61E+09 | | 8.73E+07 | | 3.97E+08 | | 9.61E+09 | |
| Dilution | Replicate 1 | Replicate 2 | Replicate 1 | Replicate 2 | Replicate 1 | Replicate 2 | Replicate 1 | Replicate 2 | Replicate 1 | Replicate 2 |
| 1:2 | | | | | | | | | | |
| 1:4 | | | | | 0.93 | 1.03 | 0.95 | 0.88 | | |
| 1:8 | 0.82 | | 0.68 | 0.66 | 1.02 | 1.18 | 1.03 | 0.86 | 0.86 | 0.79 |
| 1:16 | 0.79 | | 0.80 | 0.78 | 1.24 | 1.27 | 1.16 | 1.10 | 0.81 | 0.84 |
| 1:32 | 0.78 | 0.72 | 1.02 | 0.91 | 1.36 | 1.49 | 1.12 | 1.03 | 1.02 | 1.09 |
| 1:64 | 0.75 | 0.83 | | | | | 1.23 | | | |
| 1:128 | 0.69 | 0.74 | | | | | | | | |
| 1:256 | 0.78 | 0.72 | | | | | | | | |
| Average r ± SD** | **0.76 ± 0.05** | | **0.81 ± 0.14** | | **1.19 ± 0.19** | | **0.90 ± 0.12** | | **1.04 ± 0.13** | |
| Doubling Time+ | **31.50** | | **29.77** | | **20.15** | | **23.10** | | **26.67** | |

*Carrying capacity values correspond to the average highest value of GC/mL that could be sustained in a steady state for each isolate.

**The r value represents the replication rate measure for each data set.

+The Doubling Time represents the number of hours it takes each isolate to double.

†Greyed-out rows represent sample sets excluded from downstream analyses according to inclusion and exclusion criteria.
